# Detecting Circular RNA from High-throughput Sequence Data with de Bruijn Graph

**DOI:** 10.1101/509422

**Authors:** Xin Li, Yufeng Wu

## Abstract

Circular RNA is a type of non-coding RNA, which has a circular structure. Many circular RNAs are stable and contain exons, but are not translated into proteins. Circular RNA has important functions in gene regulation and plays an important role in some human diseases. Several biological methods, such as RNase R treatment, have been developed to identify circular RNA. Multiple bioinformatics tools have also been developed for circular RNA detection with high-throughput sequence data. In this paper, we present circDBG, a new method for circular RNA detection with de Bruijn graph. We conduct various experiments to evaluate the performance of CircDBG based on both simulated and real data. Our results show that CircDBG finds more reliable circRNAs with low bias, has more efficiency in running time, and performs better in balancing accuracy and sensitivity than existing methods. As a byproduct, we also introduce a new method to classify circular RNAs based on reads alignment. Finally, we report a potential chimeric circular RNA that is found by CircDBG based on real sequence data. CircDBG can be downloaded from https://github.com/lxwgcool/CircDBG.

## 1 Introduction

Circular RNA, the RNA in a circular form through a usually 5 to 3 phosphodiester bond, is a type of non-coding RNA [1]. Circular RNA (or circRNA) is recently recognized as a new class of functional molecule. CircRNA consists no 5 or 3 free terminus, as illustrated in Figure 1, which makes it much more stable in the cells than linear RNA [1]. CircRNAs were originally thought as the byproduct from the process of mis-splicing, and considered to be of low abundance. Recently, however, the importance of circRNAs in gene regulation and their biological functions in some human diseases have started to be recognized [1, 2, 3]. Many of these circRNAs are stable and contain exons, but they are not translated into proteins [4].

**Fig. 1.**
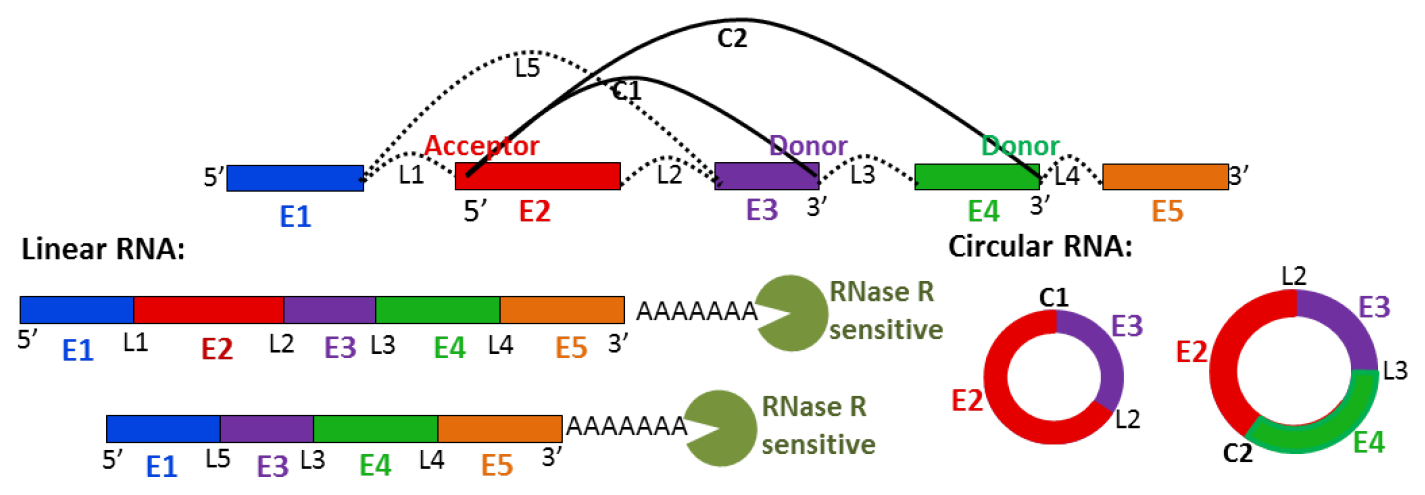
*E_i_*: exons, *L*_*i*_: linear splicing junction (for linear RNA). *C*_*i*_: circular splicing junction (for circRNA). Two sets of isoforms of linear RNA and circRNA are shown on left and right sides. 3’ in circular splicing junction is donor, while 5’ is acceptor. All linear isoforms are sensitive to RNase R. Circular isoforms show no significant decrease in abundance after RNase R treatment.

There are two types of experimental methods currently that can be used to identify circular RNA [5]. First, since circRNAs lack a poly(A) tail [6], they can be retained in rRNA-depleted libraries by using expected depletion profile to assess results. Second, circRNA can be enriched in libraries treated with RNase R to digest linear RNA and make it easier to detect lowly expressed circRNA.

With the high-throughput sequencing technologies, multiple bioinformatics tools have been developed recently for circRNAs detection from RNA sequence reads. Some of them require gene annotation while others do not. Those methods could be divided into two categories: (a) reads-mapping-based methods, such as CIRI [7]/CIRI2 [8], CIRCExplorer [9], Find-circ [10] and CircRNAFinder [11], and (b) k-mer-based methods, such as CircMarker [12].

Reads-mapping-based methods first map the RNA-seq reads onto a reference. For this purpose, CIRI uses BWA, while bowtie and Tophat (TopHat-fusion) are used by find-circ and CIRCExplorer respectively. Since BWA and bowtie do not require annotations, all RNA reads need to be mapped to the entire reference genome. As CIRCExplorer, CircRNAFinder only performs reads mapping by using the STAR in the range of annotated genomes as provided by annotation file. Those mapping methods have two major issues. First, reads-mapping-based tools is often computationally inefficient because mapping all reads can be slow, yet we note that many RNA-seq reads are irrelevant to circRNA detection. Second, these tools may miss circRNA in some cases due to errors in reads mapping [12].

Recently, we developed a k-mer-based tool called CircMarker [12], which uses an efficient k-mer table for circular RNA detection. Compared with the reads-mapping-based method, CircMarker has two major advantages. First, CircMarker looks for the circRNA-related reads for detection and does not depend on any third party mapping tool. Thus CircMarker is much faster than reads-mapping-based methods, especially for small data. Second, since the minimum comparison unit for CircMarker is a k-mer rather than reads, it can tolerant more errors and find more circular RNAs. However, CircMarker still has some issues. A key issue for CircMarker is the potential loss of information. CircMarker considers k-mers individually. That is, CircMarker does not consider the order of k-mers from either reads or exons, and this may lead to false positives when there are repetitive k-mers. Moreover, CircMarker becomes slow for large data.

In this paper, we present a new method named CircDBG for circular RNA detection with de Bruijn graph. Through experiments based on simulated and real data, we demonstrate that this new method finds more reliable circRNA with low bias, runs faster and has better performance in balancing accuracy and sensitivity than existing methods.

Finally, we introduce a new method of classifying circular RNAs based on reads alignment and report a potential chimeric circular RNA that is found by CircDBG based on real sequence data.

## 2 Method

### 2.1 High-level approach

The key idea of CircDBG is creating a de Bruijn graph based on k-mers from the boundary parts of exons in annotated genome. As shown in Figure 2, we take advantage of this graph to find the relationship between k-mer of reads and the potential donor/acceptor exon by tracking the path in the graph for circular RNA detection. Since the path provides a stronger signal for calling the two exons involved in the back splicing than individual k-mers, CircDBG can filter out more false positives than CircMarker. This is especially true when there are duplicate k-mers in exons and/or there are errors in the reads. To make CircDBG more efficient, we also develop various techniques.

**Fig. 2.**
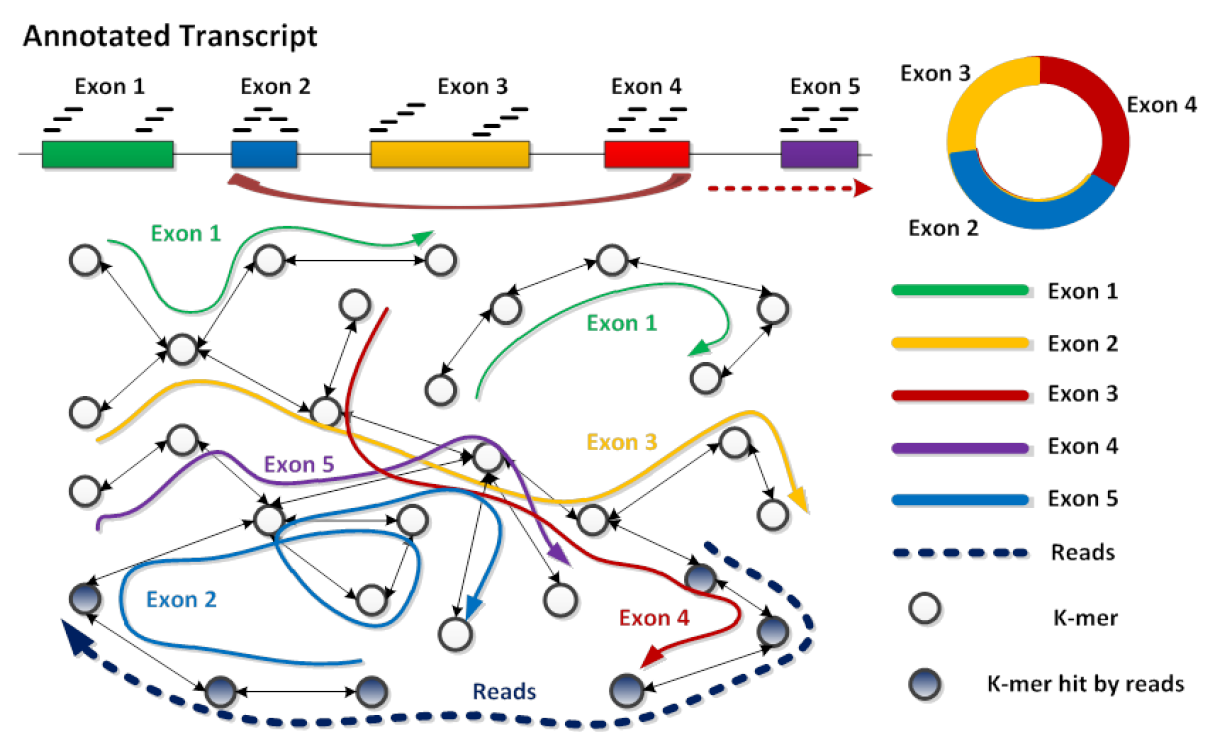
Five exons are in the genome (top left). Back splicing occurs from exon 4 to exon 2, which generates a circular RNA (top right). The bidirectional de Bruijn graph (bottom left) is built from the k-mers from each exon in the genome, where each exon is represented by the path with the same color in graph. The dotted line represents a RNA reads which supports the presence of circRNA case: the starting part of the read overlaps with the ending part of exon 4, and the ending part of the read overlaps with the starting part of exon 2.

### 2.2 CircDBG

Our new CircDBG method contains three parts: (a) building de Bruijn graph, (b) finding potential donor/acceptor sites, and (c) detecting circular RNA. First, a memory-efficient de Bruijn graph is created, which records relevant information of annotated genome. Second, we filter out circRNA-related reads and find the potential donor/accepter exon. Finally, we compare k-mers from reads with the graph for circular RNA detection. Some parameters should be determined before running CircDBG, such as the length of k-mer. The maximum length of k-mers in CircDBG is 16, and we set 15 as its default value for all data analysis reported in this paper.

#### Build de Bruijn graph

We create de Bruijn graph for each chromosome separately, and use them in parallel with RNA sequence reads for circular RNA detection. For each chromosome, only the exons that contain back splicing signal (GT-AG) are considered. The exons with length shorter than the chosen k-mer length are ignored. Since the back-splicing only occurs near the boundary of exon, and one read cannot cover the whole exon, especially when the exon is very long, we only use k-mers near the boundaries of an exon when building the graph. The length of extraction is identified as:

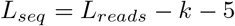

This means we require that there are at least 5 continuous k-mers from the other side of circular splicing junction. If the length of an eligible exon is shorter than 2 × *L_seq_*, the whole exon is used to build graph. All k-mers from the boundary parts of sequences are processed sequentially and converted into integers as the values of nodes in the graph. The edge of each node represents its next or previous neighbors. The procedure of creating de Bruijn graph is illustrated in Figure 3. Since the node may be shared by multiple exons or appear multiple times in one exon, multiple groups of exon information are associated with each node. Each exon information contains the nodes position and multiple links. As same as the strategy of Circ Marker, we use 5 values to represent the node position: one tag and four indexes (including chromosome, gene, transcript, and exon), which could improve the memory efficiency [12]. The tag contains 4 different values: S/E and H/T, which specifies whether the k-mer comes from the starting or ending part and whether it is close to head or tail boundary of exon respectively. Since all possible exon positions and the neighbors of each node have been recorded, this data structure can help to distinguish repetitive k-mers in the same exon or multiple exons. In addition, the closer a k-mer is to the back-splicing junction point, the higher possibility the k-mer can be contained by the supported reads. Therefore, we call the node with tag H/T as premium node. Here is an example of how we save the exon info for one node: suppose one k-mer is found in the valid part of exon 1 and exon 2, and it appears two times in exon 1. Then, the k-mer is converted into an integer and set as the key of this node. Two exon information are associated with this node: exon 1 info contains node position and one link, while exon 2 info contains node position and two links. Each link includes the key of its previous and next node.

**Fig. 3.**
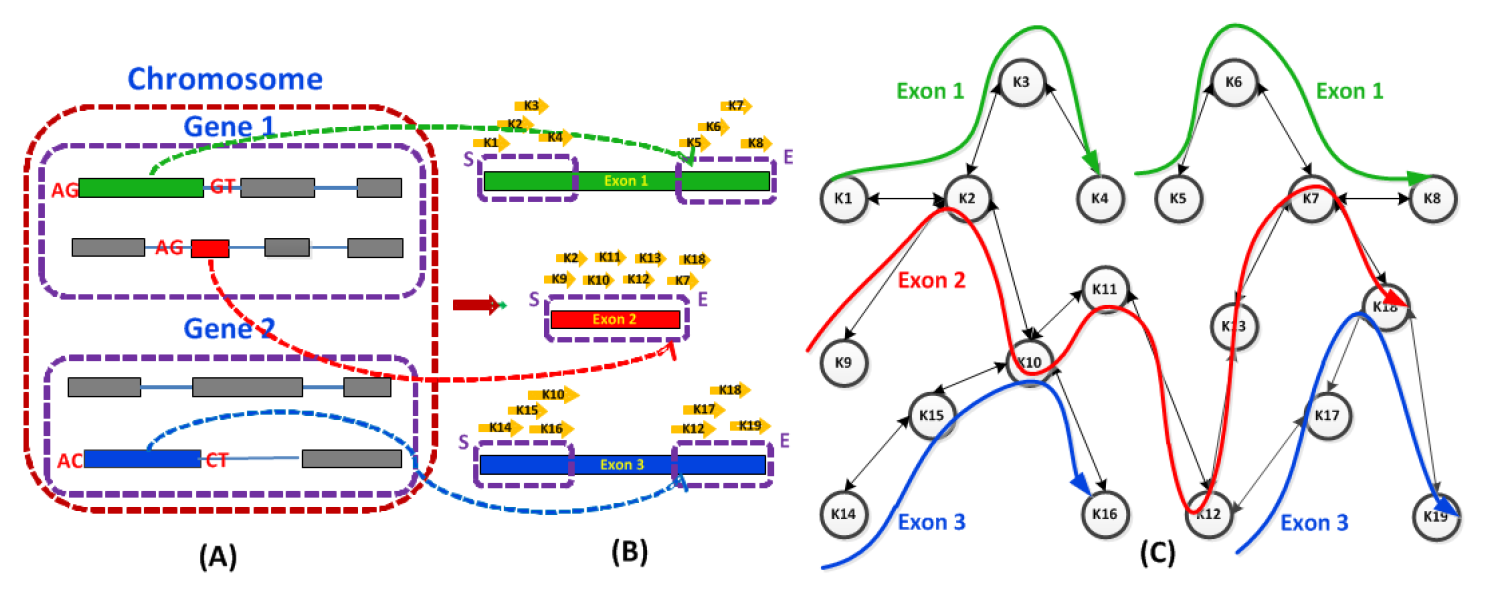
(A) Three exons with the back-splicing signal (AG-GT/AC-CT) are chosen. (B) The k-mers from the beginning part and ending part of exon 1 and exon 2 are collected, while all k-mers from exon 2 are selected since *Len*_*exon*2_ < 2 × *L_seq_*. (C) The graph is constructed by these collected k-mers: exon 1/exon 3 are represented by 2 separate green/blue paths in the graph, and exon 2 is represented by one red path.

Finally, since the last node doesn’t have the next neighbor and the first node doesn’t have the previous neighbor, we set the value of this kind of next and previous neighbor node as 0 with tag U. These two special nodes indicate the ending and beginning of the path.

#### Find potential donor/acceptor sites

We first obtain the circRNA-related reads by using the similar strategy of CircMarker: filter out reads that none of sampled k-mers from the read has matches in the graph.

In order to identify the back splicing of circRNAs, we need to search for the potential donor and acceptor sites. The donor side comes from the ending part of the exon, which is contributed by the starting part of the reads, while the acceptor side comes from the starting part of the exon, which is contributed by the ending part of the reads.

To find potential donor candidates, we sample four k-mers from the beginning to the end of the reads, and search for each k-mer’s hit in the graph. A valid hit means the k-mer can be found in the graph and it’s next neighbor in graph can be found in the read. The exon supported by at least two valid hits with tag T/E are collected as the donor candidate. For the potential acceptor candidates, we sample four k-mers from the end to the beginning of the reads, and apply the similar procedure as that of the donor candidate. There are two differences here: its previous neighbor is tracked and the valid hit should contain the tag H/S. We also collect two additional k-mers from reads for quality control. We try all combinations from the donor and acceptor candidates. If the donor and acceptor come from the same exon, we think this is the potential self-circle case. Otherwise, it belongs to regular-circle case if the donor and acceptor come from the same transcript in the order of back to front. If there are more than one candidate for each circle case, we only consider the candidate supported by the maximum number of quality control nodes. See Figure 4 for an illustration. Note that for the regular-circle candidate, if donor and acceptor nearby each other with the same sequence value, they may be from genome repeats and this candidate is ignored.

**Fig. 4.**
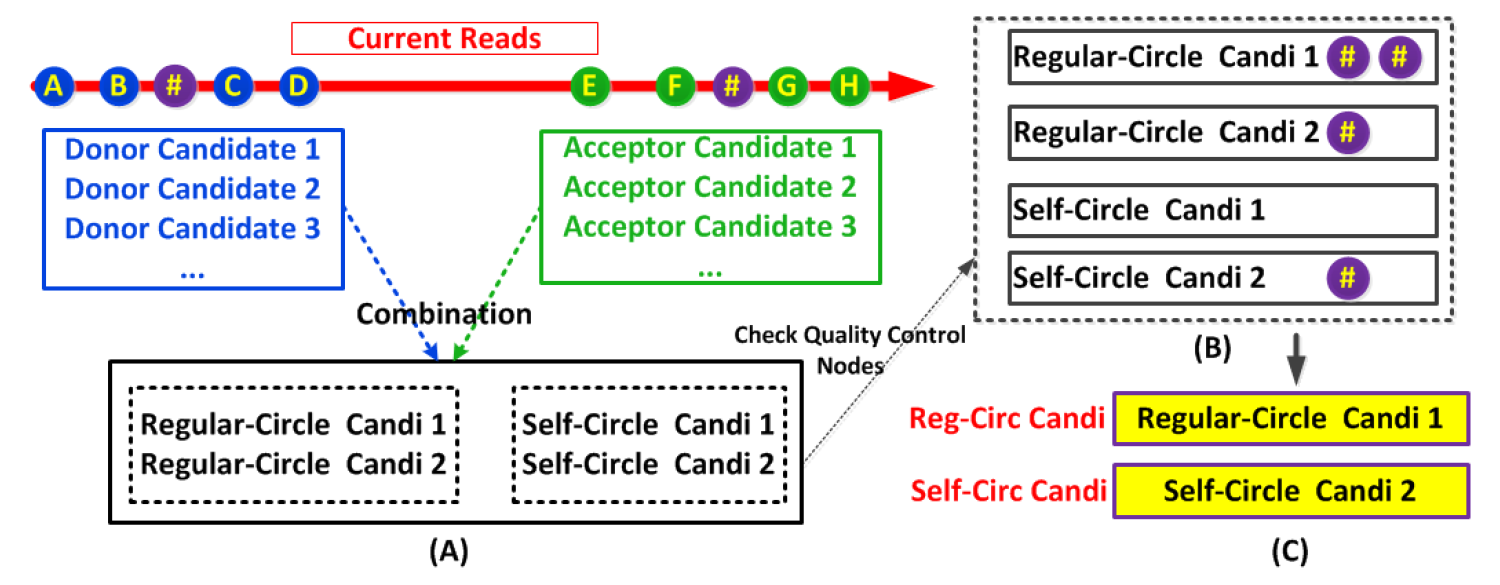
(A) Each node stands for k-mer. Four blue nodes represent the k-mers chosen from the head part of reads which are related to donor, while four green nodes represent the k-mers from the tail part which correspond to acceptor. Two purple nodes with symbol # represent the quality control nodes. Two self-circle cases and two regular-circle cases are found by the combination of donor and acceptor candidates. (B) Check whether or not quality control nodes support each circular case. (C) “Regular-Circle Candi 1” and “Self-Circle Candi 2” are kept, since they are supported by more quality control nodes than others in their case group respectively.

**Fig. 5.**
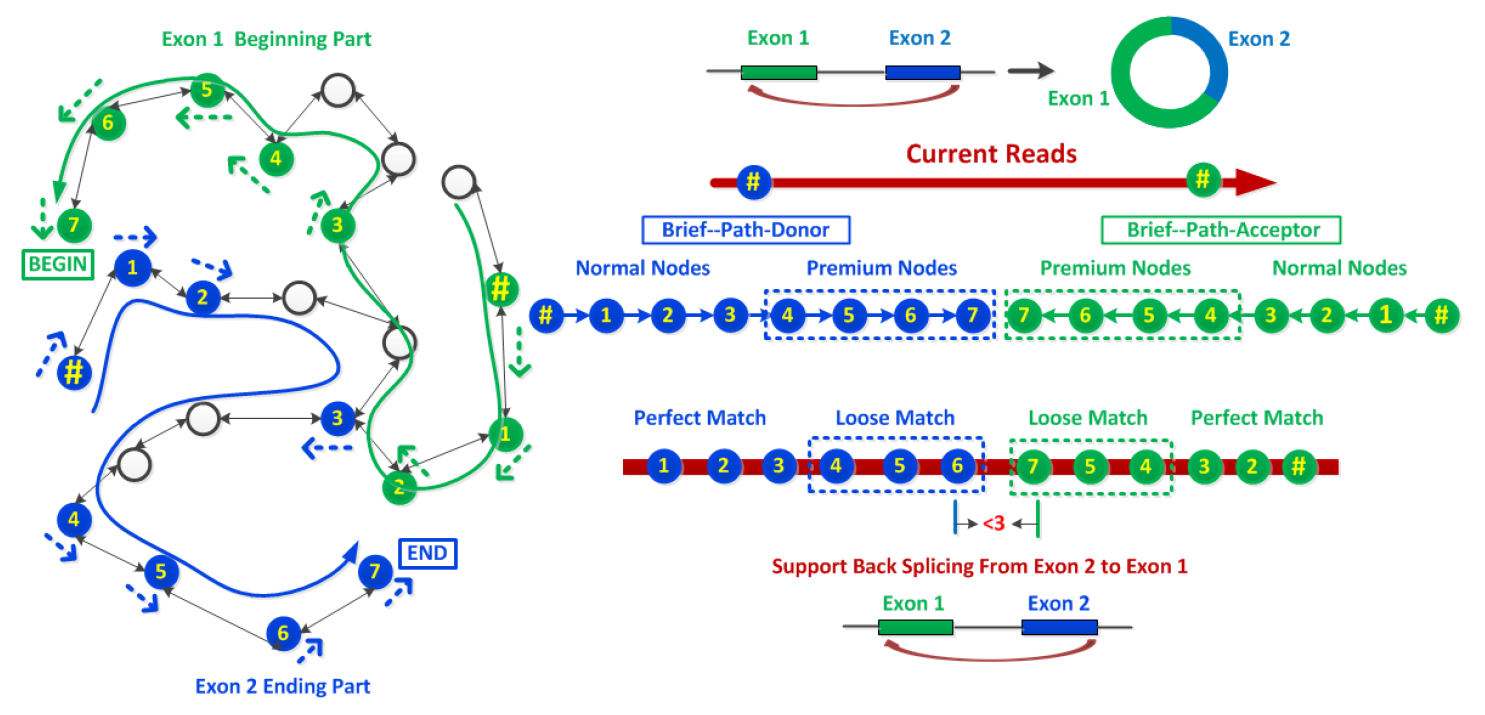
Back splicing occurs from Exon 2 to Exon 1 which generates a circle. The blue node with symbol # is the anchor node of donor and the blue path in graph represents the donor (Exon 2) by tracing backward from the anchor node. “Brief-path-donor” contains eight blue nodes in path, including four premium nodes. Green path in graph represents the acceptor (Exon 1) by tracing forward from anchor node. “Brief-path-acceptor” contains eight green nodes in path, including four premium nodes. Six nodes from the brief path of donor and acceptor can find hits in reads respectively. Since there are more than 70% hits and the distance between nodes 6 and 7 is shorter than 3 bps, this reads is considered to support the back splicing from Exon 2 to Exon 1.

#### Circular RNA detection

For each circRNA candidate, we try to find the first k-mer in the reads from the beginning to the end that can find hits in the graph with the same donor information identified in the current candidate. Then, we view this hitting node as an anchor and iteratively search for its next neighbor in the graph continuously. Once we get the path, we save a “brief-path-donor” by only keeping the first three nodes, the nodes with index divisible by 3 and the last node which contains the terminal signal in its next neighbor. This brief path can speed up the later comparison while keeping the same accuracy. Here, when we check the full path in the graph, the search is terminated if the path is longer than the length of the reads. In addition, the candidate is ignored if the length of the full path is too short. Similar procedure is applied to extract the “brief-path-acceptor” by tracing the previous neighbor continuously from the anchor node, which is the first valid hit case with the same acceptor info from the end to the beginning of the reads. The total length of these two brief paths should be long enough or contain more than two premium nodes by each of them.

Once those two brief paths are prepared, we check if the nodes in “brief-path-donor” can find hits in the reads from the beginning to end sequentially. We perform “perfect match” for regular nodes. For the premium nodes, we perform“loose match”: the node sequence is divided into three parts and is viewed as a hit if at least two parts could be matched perfectly. If the number of hits is larger than our pre-defined threshold, we consider that the donor side is well supported by current reads. Otherwise, we apply a weak threshold if the hit nodes contain at least one premium node to guarantee the donor junction point to be covered by reads. The procedure of acceptor verification is similar to that of donor case. The only difference is that we try to check if the nodes in “brief-path-acceptor” can find hits in reads from the end to beginning sequentially. Then, we check the distance between two last mapping nodes in reads from those two brief paths respectively. The circRNA candidate is kept only if the distance is < 3 bps. Finally, we merge similar candidates that share similar boundary of both donor and acceptor site with maximum 8 bps differences by using the candidate with the shortest summary length of donor and acceptor to represent the final result.

## 3 Results

Comparing different circRNA detection methods is not straightforward. The field lacks a gold standard for assessing the accuracy of their genome-wide predictions [5]. In addition, although several circular RNA databases have been released recently, such as circRNADb [13], the data in these databases come from published papers which are obtained from existing circRNA detection tools and only a few of those data have been verified through biological experiments. In this paper, we use four different strategies for evaluation. All of these strategies calculate the accuracy and sensitivity of each tool as follows, where *T* is the total called circRNAs by a tool, *T_hit_* is the number of called circRNAs which find matches in the benchmark. “Benchmark” is prepared in different ways for each strategy.

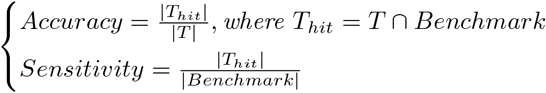

In the first strategy, we use simulated data for comparison where the simulated circRNAs are benchmark. There is no need to prepare benchmark in the second strategy, and we evaluate bias and coverage of reliable circRNAs by using public databases and real sequence data from specific tissues. In the third one, for real data, if two different datasets come from the same tissue with different experimental libraries, the intersection between the results of those two datasets could be considered as the reliable results for each tool, and the circRNAs supported by at least two tools could be viewed as the benchmark. In the last strategy, we use the intersection between the results of RNase R treated and untreated data to get the reliable results for each tool, and the circRNAs supported by at least 2 tools are viewed as benchmark. See the supplemental A for details on benchmark.

Our experiments show that no single tool always has the highest accuracy and sensitivity. Thus, we focus on comparing the balance between those two indicators by using F1 score. The F1 score is calculated by 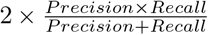. In our cases, since there is no true negatives, and all non-true positives are viewed as false positive, the precision and recall are equal to accuracy and sensitivity respectively. So the F1 score is equal to 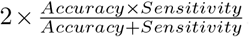. Since some tools depend on annotation files whereas some others don’t, and the majority part of back splicing comes from the exons in genome, we choose circRNAs with junction points identified by annotated genome for comparison.

Due to the space limit, the results of the second strategy are shown in the supplemental B.1: CircDBG finds more circRNAs recorded in database than other tools, and it always gets the largest coverage (20 of 23) in each chromosome respectively. Moreover, CircDBG is the best tool with the lowest bias (contain majority results of other tools) and the fastest running time. The results of other three strategies are presented below.

### 3.1 Simulated data

We use the latest simulator released by CIRI2 [8] to simulate circRNAs and RNA-seq reads. The length of simulated paired-end reads is 101 bps, and the coverages of circRNA and linear RNA are 10x and 80x respectively. The error rate is 1%. The major/minor normal distribution insertion length is 320/550, and the percentage of splicing for skipping exon is 40%. The reference and annotation file come from human chromosome 1 (GRCh37, version 18). The simulated paired-end reads contain 1,115,738 pairs. 295 circular RNAs are simulated as benchmark. Accuracy, sensitivity and F1 scores are calculated for each tool.

As shown in Figure 6, both accuracy and sensitivity of CircDBG are around 94%, and it gets the highest F1 score (0.9406). This means that CircDBG is the best tool for balancing accuracy and sensitivity. CircDBG also has the fastest running time.

**Fig. 6.**
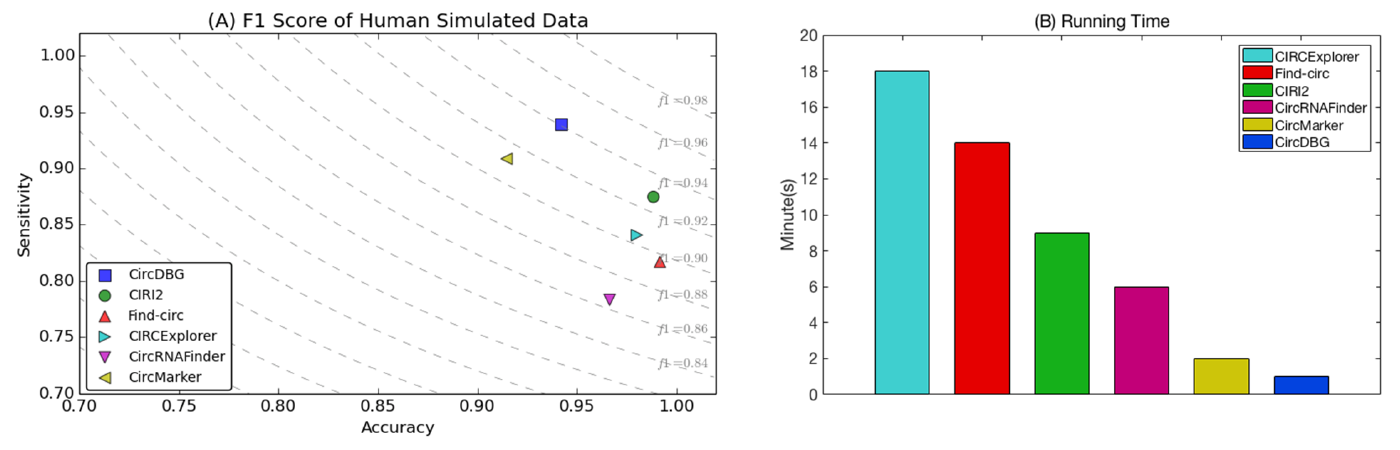
(A) Dotted line represents a fixed F1 score. The closer to top right, the higher F1 score it is. (B) Running time (in minutes).

### 3.2 Real data: two different prepared libraries from same tissue

This comparison is based on two different libraries from the same issue. Intuitively, true circRNAs should be called for both libraries. We get the reliable results for each tool by taking the intersection between two differently prepared libraries from the same tissue. The results that are supported by both libraries are viewed as reliable. RNA-seq reads SRR4095542 and SRR5133906 are used for data analysis. The first library (43,488,788 paired-end reads) is prepared by 3 glioma and paired normal brain tissue. For the second library (54,732,199 paired-end reads), ten human glioblastoma samples are mixed as tumor group, and their periphery normal tissues are mixed as control group. Total RNAs in the second library are extracted and treated with RNase R to remove the linear RNAs. The length of reads is 150 bp in both libraries. Recall that the intersection of the called circRNAs from two libraries is used as the final result for each tool, and circRNAs supported by at least 2 tools are viewed as benchmark.

Our results are shown in Figure 7. CircDBG has the highest F1 score (0.9539). In addition, the accuracy of CircDBG is 98.65% with the highest sensitivity and the fastest running time.

**Fig. 7.**
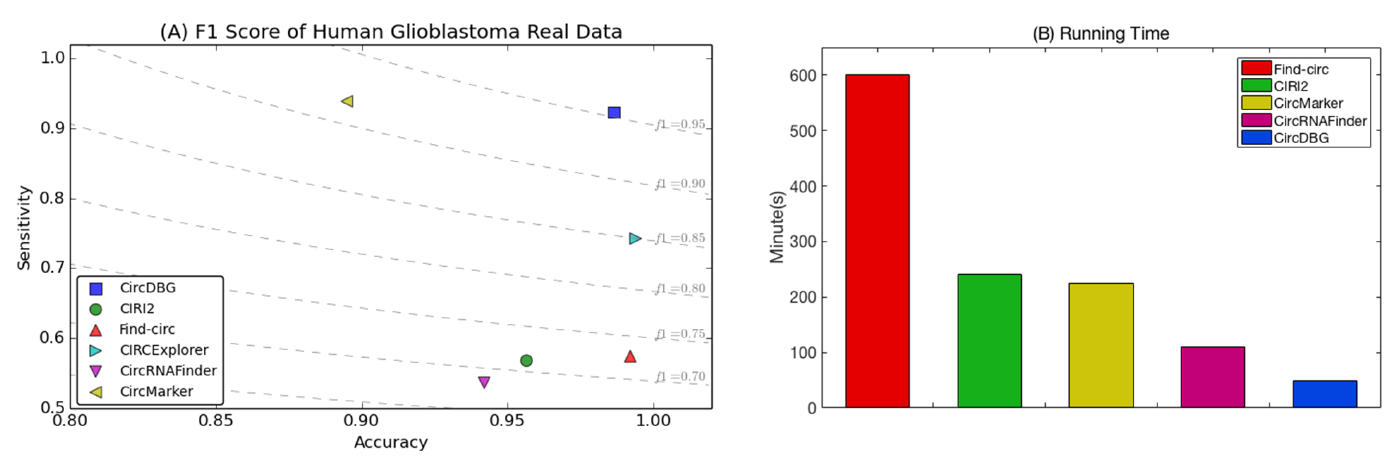
(A) Dotted line represents a fixed F1 score. The closer to top right, the higher F1 score it is. (B) CIRC Explorer takes more than 36 hrs and is not shown here.

### 3.3 Real data: RNase R treated and untreated samples

This comparison is performed with RNase R treated and untreated libraries. We collect two sets of treated and untreated reads from homo sapiens and mus musculus respectively. A circRNA is viewed as reliable if it can be found by both RNase R treated and untreated reads (linear RNAs tend to degrade by RNase R treatment). SRR1636985 (treated, 13,309,745 paired-end reads) and SRR1637089 (untreated, 44,933,450 paired-end reads) from HeLa cells are used for human. The length of reads in both libraries is 101 bps. For mouse libraries, SRR2219951 (treated, 22,330,976 paired-end reads) and SRR2185851 (untreated, 32,939,809 paired-end reads) are selected, which are prepared by mouse brain at the age of 8 to 9 weeks. The length of reads in the two groups varies with the maximum 100 bps. For each species, we obtain final results for each tool by taking the intersection between the results based on treated and untreated reads and build the benchmark by choosing the circRNAs supported by at least two tools.

As shown in Figure 8, CircDBG has the highest F1 score and the fastest running time in both human (F1 Score: 0.9589) and mus musculus (F1 Score: 0.9424). The accuracy of CircDBG in human and mouse are 98.24% and 99.25% respectively. CircDBG and CircMarker are the top two tools that get the highest sensitivity in both human and mouse libraries.

**Fig. 8.**
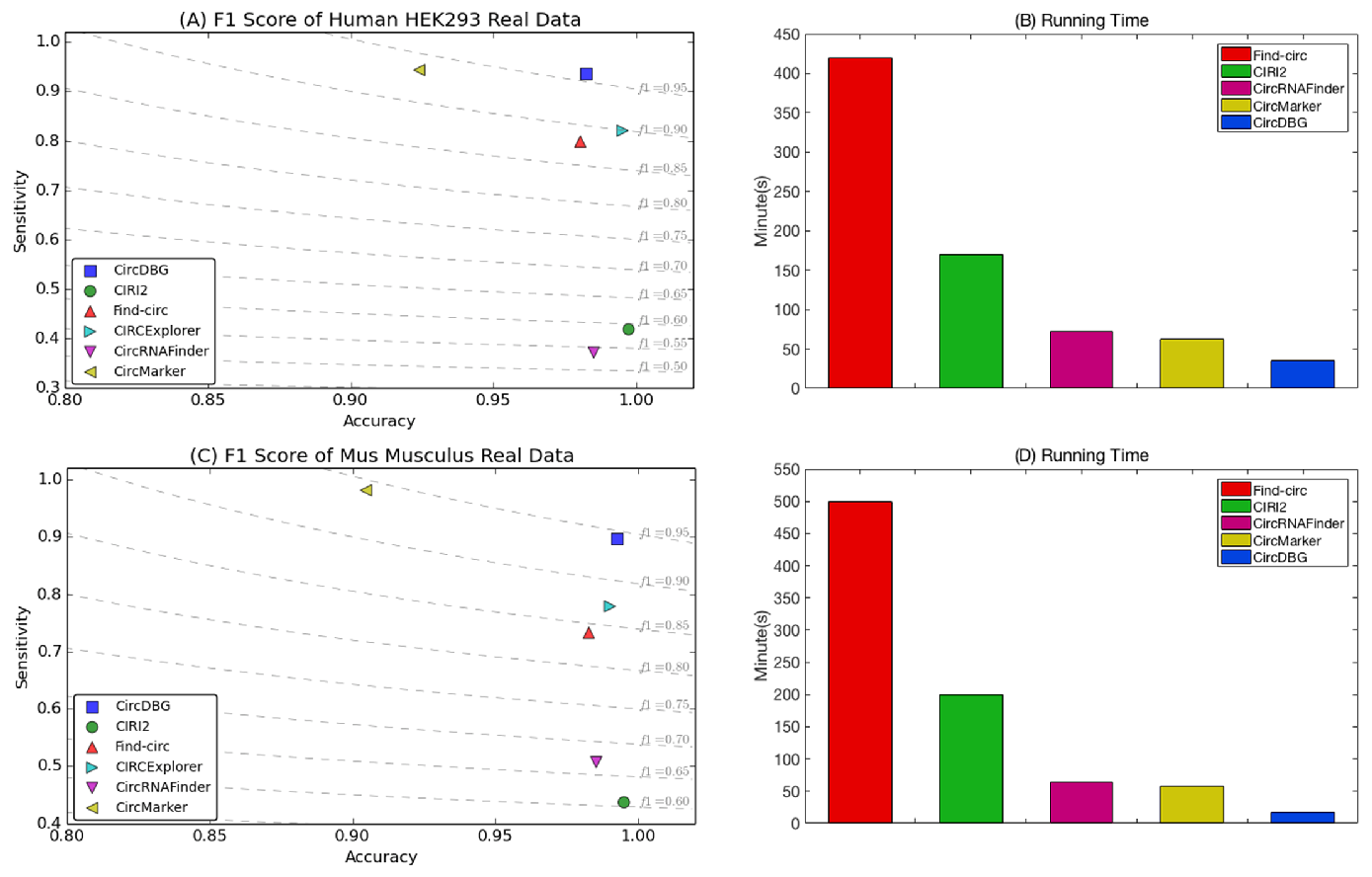
(A) and (B) are the comparison results of human data. (C) and (D) are the result of mouse. CIRCExplorer takes 15 hrs and 21 hrs for human and mouse respectively, and is not shown in (B) and (D).

## 4 Case study: chimeric circular RNA

We develop a novel evaluation scheme, which analyzes what reads contribute to the calling of circRNAs. In this scheme, CircRNA reference is generated by linking the ending part of donor exon with starting part of acceptor, and the circRNAs detected by CircDBG are classified into 5 categories based on the alignment result of reads. See the supplemental C for details.

We notice that there is a special case when classifying the detected circRNA with real data: some reads not only support regular-circle case, but also support the self-circle case for the exon contained by current regular-circle case, which is illustrated in Figure 9. This case may relate to chimeric phenomenon in circular RNA which can be comprehended as “circle in circle”.

**Fig. 9.**
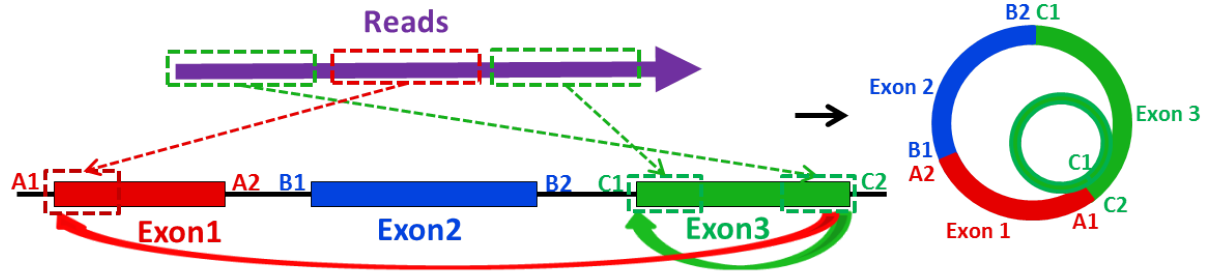
The first and the second part of reads support the regular-circle RNA case from exon 3 back to exon 1, while the first and the third part of reads support the self-circle RNA case of exon 3.

We collect this type of chimeric circular RNA in real data and find this phenomenon exists in all real data we analyze. For example, in H9 hESCs, 102 chimeric cases can be found among the total 11,931 circular RNA. Although the percentage seems to be small, the number of the chimeric cases is significant.

## 5 Conclusion

In this paper, we develop a new method called CircDBG for circular RNA detection, which is based on de Bruijn graph. The graph represents the relationship between k-mers in original exon and reads. This contributes to more accurate results compared with the existing k-mer-based methods. We conduct extensive experiments and demonstrate CircDBG outperforms existing tools, especially in saving running time, reducing bias and improving capability of balancing accuracy and sensitivity. CircDBG is the stand alone tool and does not depend on any other third party tools. CircDBG can be downloaded from: https://github.com/lxwgcool/CircDBG.

## Acknowledgement

This work is supported by grants IIS-1447711 and IIS-1526415 from US National Science Foundation to YW.

## Supplemental materials

### A Benchmark used for comparison

The benchmark is represented by the formula below, where *T* is the final result of current tool, which comes from the intersection between the detection results of dataset A (e.g. treated) and B (e.g. untreated). “n” is the total number of tools, and the final benchmark contains all the results which are supported by at least two tools.

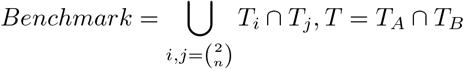

### B Additional results

#### B.1 Real data: circRNADb with tissue H9 hESCs

We choose the database circRNADb in this comparison, and all records in database are viewed as reliable circRNA, since these records were collected from several published studies and only the circRNAs supported by more than one read are recorded [13]. There are two goals of this comparison. First, we want to examine how well the public database is supported by each tool. The larger coverage in database the results from a tool has, the better support the tool is. Second, we evaluate the bias of each tool by checking the overlap between the results of the current tool and others respectively. The larger overlap means the lower bias. All circRNAs recorded in circRNdb come from Homo Sapiens, and there are total 10,631 circular RNAs from H9 hESCs, which are used for comparison. The real reads SRR901967 are chosen for data analysis. These reads are specially designed for examining circular RNAs in H9 human embryonic stem cells with RNase R treated. It contains 41,342,095 single reads with the length 100 bps. We use three statistics in the comparison, including the number of circular RNAs hitting database (including the hitting situation in each chromosome respectively), the overlap between the results of each tool, and the running time.

Our results are shown in Figure S1. CircDBG covers more circular RNAs recorded in database than other tools, and it always gets the largest coverage (20 of 23) in each chromosome respectively. In addition, CircDBG and CircMarker overlap with more results from other tools. Moreover, CircDBG performs better than CircMarker in overlapping with two tools (CIRI2 and CIRCExplorer) and similarly in overlapping with other two tools (Find-circ and CircRNAFinder), which means CircDBG is the best tool with the lowest bias. Finally, CircDBG is much faster than other tools.

**Fig. S1.**
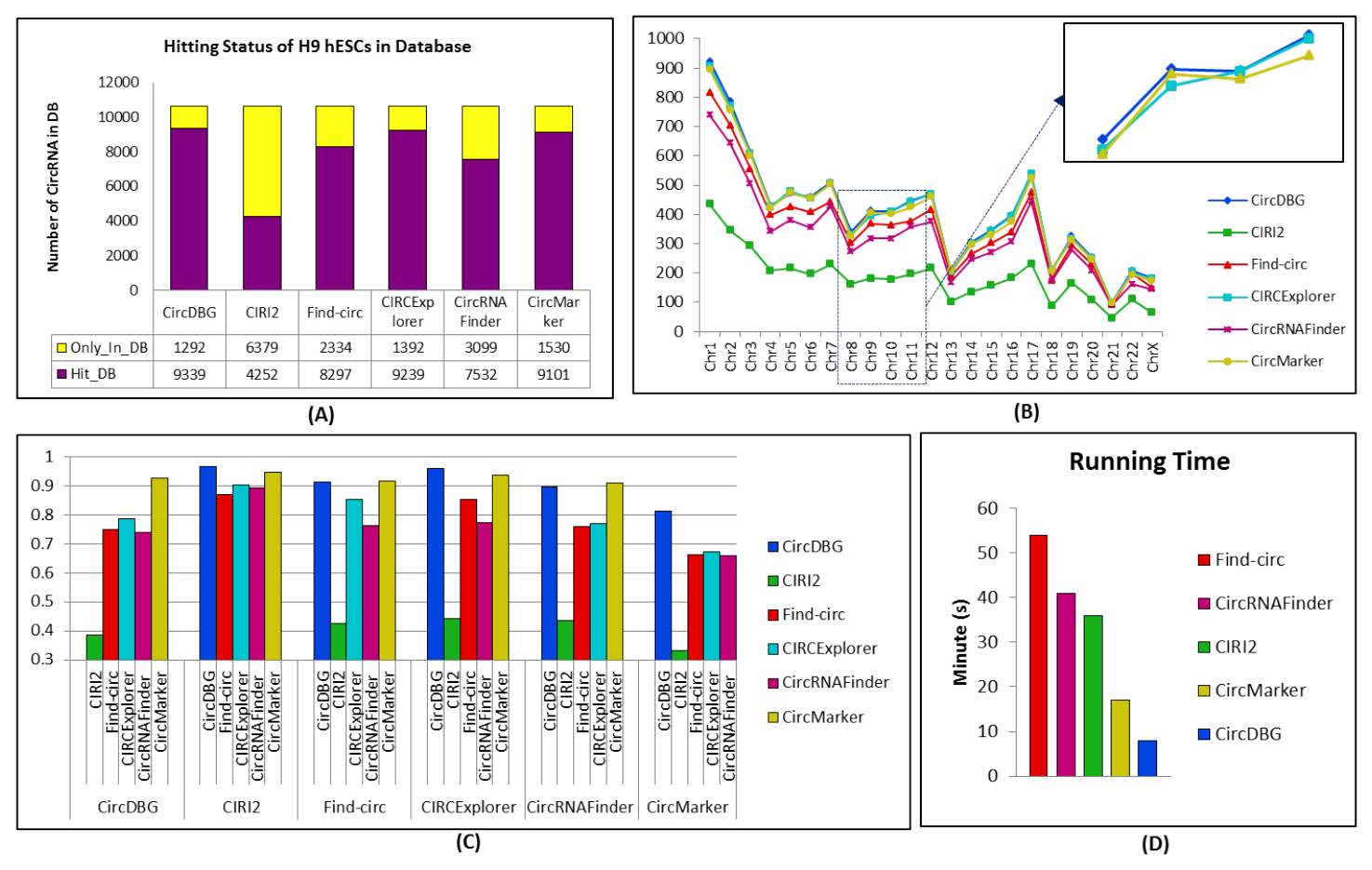
(A) The yellow bar represents the circRNA from H9 hESCs recorded in database but not contained by the results of current tool, while the purple bar means the circRNA contained by both of them. (B) The plots represent circRNA contained by the current tool and database in each chromosome. (C) There are 6 groups, and the bars in each group mean the number of results of current tool covered by other tools. (D) CIRCExplorer takes more than 15h and is not shown here.

### C Classification of circRNA by reads

We note that there are some differences among the results of different tools. In this section, we take a closer look at the results in order to see if there are some specific groups of circRNAs that can only be found by CircDBG.

Since each circular back splicing is detected by reads, we want to find out how those reads contribute to circRNA detection. Intuitively, if the error rate of reads is high, this reads may not be used for detection by some tools. In addition, if the part of a read which supports donor or acceptor is too short, the read may be ignored by some tools as well. Moreover, if one part of reads doesn’t match either donor or accepter, it may also be discarded by some tools. Now, we align circRNA with their supported reads by BLAST. Five different categories are identified, including LowQuality, Imbalance, AdditionalPart, Bad and Good. “LowQuality” means the lowest error ratio is larger than 1% for all supported reads. “Imbalance” means the longest alignment segment of either the donor part or the acceptor part is shorter than 25 bps. If the minimum difference between alignment and reads length is larger than 10 bps, it is considered to be “AddionalPart”. “Bad” means the junction point is out of the alignment range for all supported reads. Otherwise, the quality of current circRNA is set as “Good”. All of these five categories are illustrated in Figure S2.

**Fig. S2.**
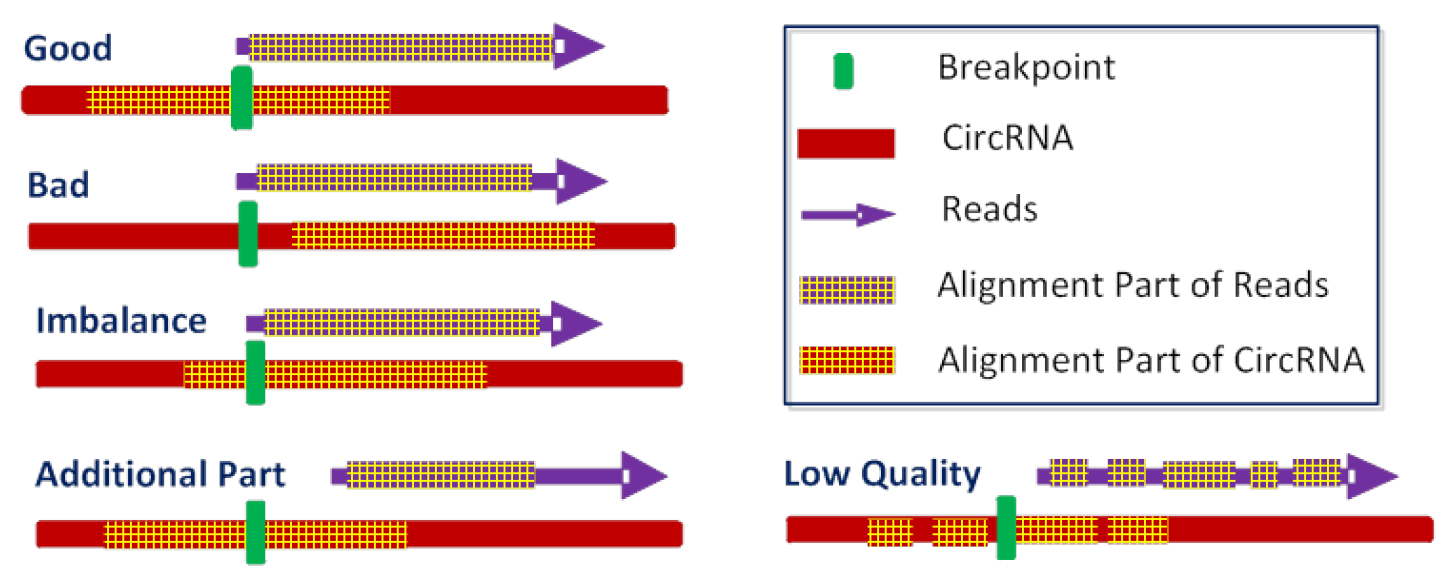
circRNA reference (Red Bar) is generated by linking the ending part of donor exon with starting part of acceptor, and green bar is used to represent the circular junction point. “Good” means the most part of reads can be aligned to circRNA, and the length of alignment part in either side of junction point is > 25 bps. “Bad” means the junction point is out of alignment part in circRNA. “Imbalance” means the alignment part on one side of junction point is too short. “Additional Part” means a continuous part of reads fails to align to circRNA reference. “Low Quality” means some gaps appear in alignment results.

Since CircDBG outputs the support reads of each detected circRNA, we apply this classification strategy to the called circRNAs by CircDBG in the real data. We find circRNA in “Good” category could be detected by most existing tools. However, the majority of circRNAs in “Imbalanced”, “Additional part” and “Low Quality” categories are only detected by CircDBG and CircMarker. For example, in H9 hESCs, the detected number of circRNA for the categories of “Good”, “Bad”, “Additional Part”, “Imbalance” and “LowQuality” are 8,604, 1, 654, 1, 2,532 and 139, and less than 30% of “Additional Part”, “Imbalance” and “LowQuality” can be found in the results of reads-mapping-based methods. The tool named “CircAssistant” is developed to make this classification and to report the chimeric circular case based on the result of CircDBG. It could be downloaded from https://github.com/lxwgcool/CircDBG/Cir_Assistant.

